# Dynamic expression of risk genes for schizophrenia and bipolar disorder across development

**DOI:** 10.1101/333104

**Authors:** Nicholas E Clifton, Eilís Hannon, Arianna Di Florio, Kerrie L Thomas, Peter A Holmans, James TR Walters, Michael C O’Donovan, Michael J Owen, Andrew J Pocklington, Jeremy Hall

## Abstract

Common genetic variation contributes a substantial proportion of risk for both schizophrenia and bipolar disorder. Furthermore, there is evidence of significant, but not complete, overlap in genetic risk between schizophrenia and bipolar disorder. It has been hypothesised that genetic variants conferring risk for these disorders do so by influencing brain development, leading to the later emergence of symptoms. The comparative profile of risk gene expression for schizophrenia and bipolar disorder across development over different brain regions however remains unclear. Using genotypes derived from genome wide associations studies of the largest available cohorts of patients and control subjects, we investigated whether genes enriched for schizophrenia and bipolar disorder association show a bias for expression across any of 13 developmental stages in prefrontal cortical and subcortical brain regions. We show that genes associated with schizophrenia have a strong bias towards increased expression in the prefrontal cortex during early midfetal development and early infancy, and decreased expression during late childhood which normalises in adolescence. Risk-associated genes for bipolar disorder shared this postnatal expression profile but did not exhibit a bias towards expression at any prenatal stage. These results emphasise the dynamic expression of genes harbouring risk for schizophrenia and bipolar disorder across prefrontal cortex development and support the view that prenatal neurodevelopmental events are more strongly associated with schizophrenia than bipolar disorder.

## INTRODUCTION

Schizophrenia and bipolar disorder are disorders of mainly adult onset, yet both have been considered to have earlier developmental origins^1–4^. In the case of schizophrenia, one influential hypothesis has been that events occurring during prenatal brain development confer risk for the disorder which remain latent and are only expressed later as symptoms in the context of brain maturation^1,5^. Prefrontal cortical regions, which don’t fully mature until the late second decade of life, have been considered one potential substrate of such early developmental vulnerability for later schizophrenia^1,5,6^. The importance of prenatal neurodevelopment in risk for schizophrenia is supported by epidemiological evidence showing that prenatal and perinatal events such as infection, famine and obstetric complications can increase risk for the disorder, and by the observation that children with subtle neurological, cognitive and behavioural impairments are at enhanced risk for later developing the condition^3,4,7,8^.

There is less evidence that early neurodevelopmental events play an important role in the development of bipolar disorder^4,9^. In general, fewer epidemiological studies have examined potential associations between prenatal / perinatal events such as maternal infection and obstetric complications and risk for later bipolar disorder^9^. Whilst some evidence exists for a role of prenatal influenza infection and low birth weight or prematurity in risk for bipolar disorder, results have not been consistent across studies^9–15^. Furthermore, there is not strong evidence that individuals who later develop bipolar disorder had poorer premorbid cognitive functioning and social adjustment or increased rates of subtle neurological symptoms^4,16^.

There has also been considerable interest in the potential role of developmental events occurring postnatally during childhood and adolescence in the development of adult onset psychiatric conditions such as schizophrenia and bipolar disorder^4,6,17^. It is well established that extensive neuronal maturation occurs across childhood and adolescence^6^. In particular, prefrontal cortical areas show considerable developmental change across late childhood and adolescence with extensive synaptic pruning and elimination of excitatory synapses shaping the late-maturing cortex^6,18^. There is also evidence that environmental risk factors operating across the period of childhood and adolescence can increase risk for both schizophrenia and bipolar disorder. In particular childhood abuse is associated with an increased risk of both disorders^19–21^, and maternal loss prior to the age of 5 is associated with bipolar disorder^22^. Furthermore, cannabis use has been found to have particularly pronounced effects on risk for the development of psychosis if exposure occurs prior to or during the adolescent period^6,23,24^.

Both schizophrenia and bipolar disorder are known to have a high degree of heritability^25,26^. Recent large scale genomic studies have made major progress in determining the genetic architecture of schizophrenia and bipolar disorder^27–29^. Genetic risk for both conditions is highly polygenic in nature with a major contribution from a large number of common variants, which collectively account for a significant proportion of heritability for these conditions^30,31^. There is considerable, although not complete, overlap seen between common variants conferring risk for schizophrenia and bipolar disorder as identified in genome-wide association studies^32,33^. However rarer but more penetrant genomic risk variants, such as copy number variants, are generally found to be more enriched in schizophrenia than in bipolar disorder^34,35^. These findings have contributed to the development of a “neurodevelopmental continuum” model in which schizophrenia is considered to be associated with a greater load of early neurodevelopmental insults than bipolar disorder^16,34,35^.

Previous studies examining the expression of the top genome-wide association study (GWAS) common variant risk-associated loci for schizophrenia in post-mortem tissue have confirmed a high level of expression in pre-natal brain development, consistent with the view that risk for schizophrenia may have early neurodevelopmental origins^5,36–38^. However, less is known about the early expression of bipolar-associated genes, or the comparative profile of risk gene expression for schizophrenia and bipolar disorder across development and into adulthood. In this study we therefore used data from large scale GWAS of schizophrenia and bipolar disorder combined with gene expression data across development from the BrainSpan dataset^39^, to examine the dynamic expression of genes harbouring common risk variants across developmental stages from prenatal life to adulthood.

## METHODS

### Neurodevelopmental transcriptome

Transcriptomic data was obtained from the Allen Institute BrainSpan Atlas:^39^ a resource providing RNA sequencing-derived gene expression data of post-mortem brain tissue from 38 individuals between 8 post-conceptual weeks and 40 years of age (Supplementary Table 1). The data was filtered to remove samples with low RNA integrity (RIN<7) and genes which were unexpressed (zero reads per kilobase per million (RPKM) in all samples) or did not possess an NCBI/Entrez gene ID, and any duplicate entries. RPKM gene expression values were adjusted to control for RNA integrity, ethnicity and gender by fitting a linear regression model of expression with these factors as independent variables; the residuals and intercepts from this model were used as the adjusted expression values. Subjects were grouped into 13 developmental stages (Supplementary Table 1).

### Calculation of gene expression scores

For each developmental stage, each gene was assigned a relative expression score reflecting the degree of expression relative to all other developmental stages. Expression scores were calculated by fitting a linear regression model for each gene and developmental stage: expression = brain region + developmental stage, where expression is the adjusted expression value of the gene and developmental stage is a binary variable denoting whether the sample was from the stage or not. This model was fitted to selected groups of brain regions. The brain region covariate was included to account for differences in gene expression between subregions. We defined the expression score = R^2^ ×; sgn(coefficient), so that for any developmental stage genes with high expression relative to other stages will have large positive scores, those with low expression relative to other stages will have large negative scores, and those whose expression is relatively constant across development will have scores close to zero. We structured analyses in a hierarchical fashion whereby developmental expression was first analysed using the combined data from all brain regions; a second tier of analyses was then performed on major groups of subregions.

### Genotype data

Schizophrenia and bipolar disorder single nucleotide polymorphism (SNP) association summary statistics were taken from previously described GWAS meta-analyses^27,28^. A combined schizophrenia sample of 40 675 cases and 64 643 control subjects consisted of 11 260 cases and 24 542 controls from studies of UK patients with schizophrenia taking clozapine (CLOZUK)^27^, and 29 415 cases and 40 101 controls from a large-scale GWAS performed by the Psychiatric Genomics Consortium (PGC)^29^. The bipolar disorder sample comprised of 20 352 cases and 31 358 controls from 32 cohorts of European descent, compiled as part of a recent PGC GWAS^28^.

Schizophrenia and bipolar disorder GWAS SNPs were filtered to include only those with a minor allele frequency ≥ 0.01 and imputed INFO score ≥ 0.6. Additional filters were applied to exclude the extended major histocompatibility complex (xMHC) region and the X chromosome, consistent with previous analyses^27^.

### Gene property analysis

The relationship between gene expression scores and enrichment for association with schizophrenia or bipolar disorder was determined for each developmental stage using gene property analyses in MAGMA v1.06^40^. Briefly, common SNP association *P*-values were combined into gene-wide *P*-values (*via* the MAGMA SNP-wise mean model), using a window of 35 kb upstream and 10 kb downstream^41^ of each gene in order to include SNPs within regulatory regions. Only protein-coding genes were included in the analysis. The gene property analysis method performs a linear regression of gene-wide association against a gene-level property (here, relative expression score), in which covariates are included to correct for potential confounds. Our analyses were two-tailed and included correction for gene size and SNP density. In conditional analyses, gene expression scores for the conditioned developmental stage were included as covariates. At each tier of analysis, *P*-values were Bonferroni adjusted for the number of developmental stages or brain regions analysed.

## RESULTS

### Whole brain developmental expression profile of risk genes for schizophrenia and bipolar disorder

We used gene property analysis in MAGMA^40^ to determine whether gene expression during particular developmental periods is correlated with enrichment for common variant association for schizophrenia or bipolar disorder. In a combined analysis of gene expression data from all brain regions, we observed a non-uniform relationship between relative gene expression and enrichment for schizophrenia association across development (**Figure 1a**). Genes mediating risk to schizophrenia tended to be more highly expressed during the Early Midfetal 2 developmental stage (adjusted *P*=0.014). Conversely, an inverse relationship between gene expression and schizophrenia risk was observed at Late Childhood (adjusted *P*=0.038), implying low expression of risk genes, whilst relative expression at other stages was non-significant.

**Figure 1.**
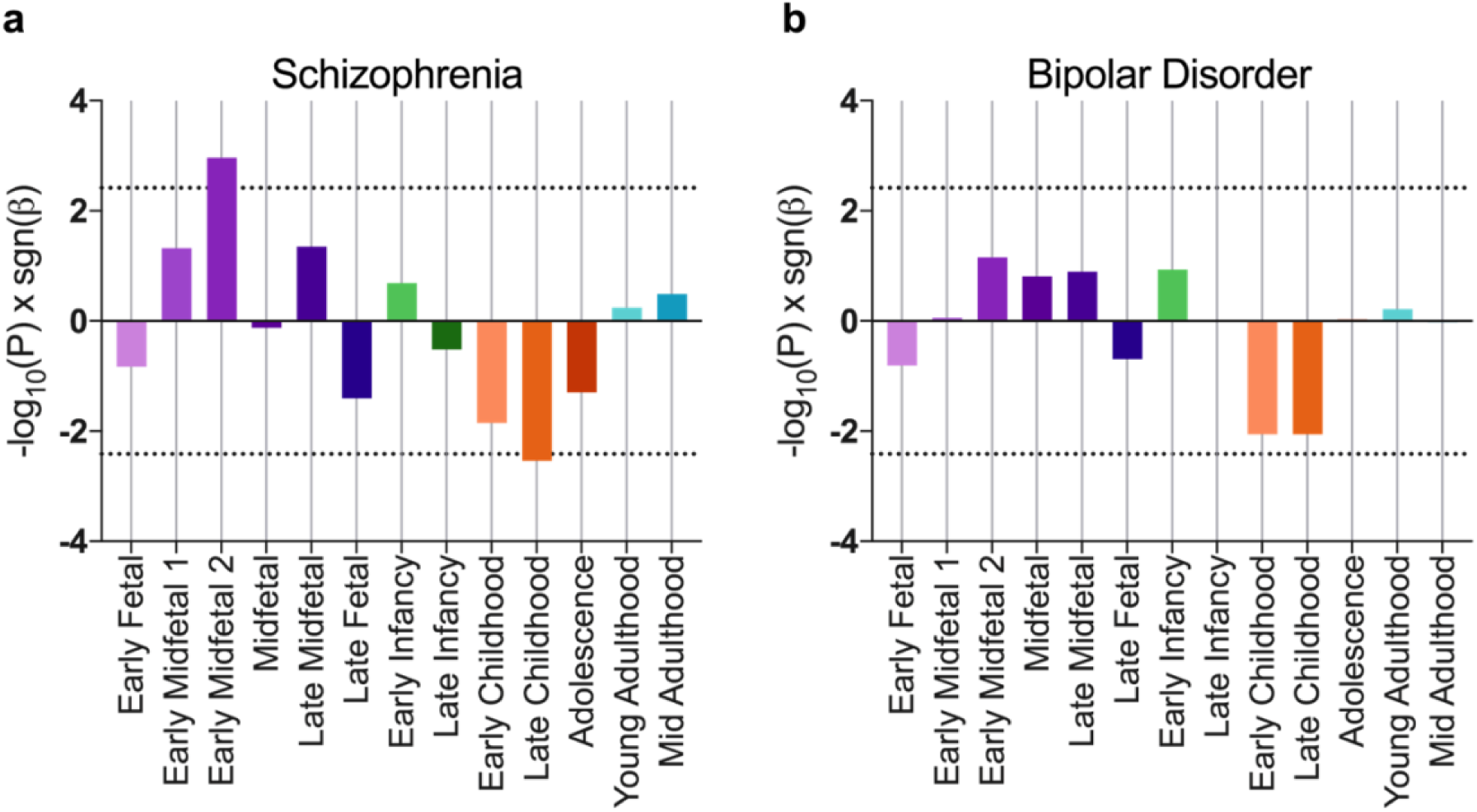
Neurodevelopmental expression bias of genes mediating risk to schizophrenia (**a**) and bipolar disorder (**b**). Gene property analysis against case-control common variant data was performed on gene expression scores derived from RNA sequencing data of all brain regions. Bars represent -log_10_(*P*) ×;sgn(β), where *P* is uncorrected following gene property analysis and β is the corresponding regression coefficient. Dotted lines show significance thresholds accounting for multiple testing across 13 developmental stages.

Brain-wide analyses of bipolar disorder GWAS data showed some variation of risk gene expression bias, although no significant relationship between gene expression and gene-wide association for bipolar disorder risk was observed at any single developmental stage (**Figure 1b**).

### Cortical and subcortical analyses of risk gene expression across development

Since trajectories of neurodevelopmental gene expression vary considerably between brain regions^39^, we sought to determine whether relationships with risk for schizophrenia or bipolar disorder were more pronounced for specific subregions of the brain. We tested these relationships in grouped sets of prefrontal cortex (medial prefrontal cortex, dorsolateral prefrontal cortex, ventrolateral prefrontal cortex and orbital frontal cortex), non-prefrontal cortex (posterior superior temporal cortex, inferolateral temporal cortex, primary auditory cortex, primary visual cortex, primary motor cortex, posteroventral parietal cortex and primary somatosensory cortex) and subcortical regions (amygdaloid complex, thalamus, striatum and hippocampus).

For schizophrenia, we observed increased expression of associated genes in prefrontal cortex at Early Midfetal 1 (adjusted *P*=6.3×10^−4^) and Early Infancy (adjusted *P*=0.034; **Figure 2a**). Gene expression at Early Midfetal 2 development was nominally related to schizophrenia risk (adjusted *P* = 0.053). Consistent with whole brain analyses, decreased gene expression in prefrontal cortex at Late Childhood was strongly correlated with schizophrenia risk (adjusted *P*=1.7×10^−7^). In non-prefrontal cortex and subcortical regions, gene expression was not correlated with schizophrenia association at any developmental stage (**Figure 2c,e**).

**Figure 2.**
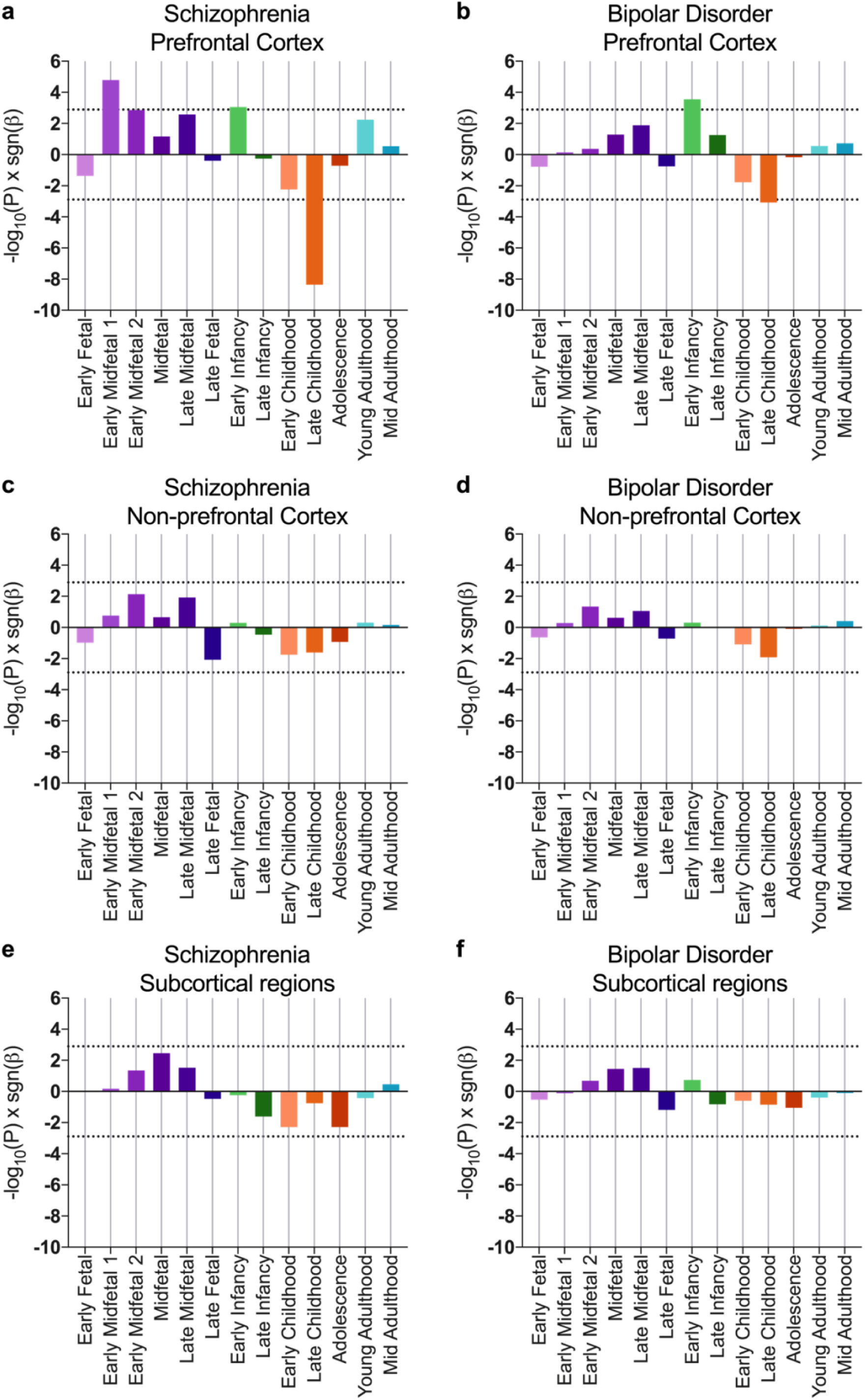
Cortical and subcortical relationships between gene expression and risk for schizophrenia (**a**,**c**,**e**) or bipolar disorder (**b**,**d,f**). Prefrontal Cortex includes medial prefrontal cortex, dorsolateral prefrontal cortex, ventrolateral prefrontal cortex and orbital frontal cortex. Non-prefrontal cortex includes posterior superior temporal cortex, inferolateral temporal cortex, primary auditory cortex, primary visual cortex, primary motor cortex, posteroventral parietal cortex and primary somatosensory cortex. Subcortical regions include amygdaloid complex, thalamus, striatum and hippocampus. Bars represent –log_10_ (*P*)×; sgn(β) in gene property analyses, where *P* is uncorrected following gene property analysis and β is the corresponding regression coefficient. Dotted lines show significance thresholds accounting for multiple testing for 13 developmental stages and 3 subregion groups.

Subregion analysis of the expression of bipolar associated genes revealed that genetic association with bipolar disorder was correlated with high prefrontal cortex expression during Early Infancy (adjusted *P*=0.011), and low prefrontal cortex expression during Late Childhood (adjusted *P*=0.033; **Figure 2b**). Whilst these postnatal expression profiles are akin to those observed for genes contributing risk to schizophrenia, prenatal expression did not correlate with risk for bipolar disorder. Gene expression in non-prefrontal cortex or subcortical regions did not correlate with bipolar disorder risk at any developmental stage (**Figure 2d,f**).

### Investigation of overlapping signals between developmental stages

We next investigated whether genes mediating the relationship between common variant association and prefrontal cortex expression during early developmental stages were also responsible for the relationship found in Late Childhood. To determine whether these signals are driven by overlapping sets of genes, we performed conditional gene property analyses for the relevant developmental stages.

For schizophrenia, conditioning on Late Childhood expression resulted in reduced correlation of prefrontal cortex expression at Early Midfetal 1 with common variant association (uncorrected *P*=0.0054). This indicates that a substantial proportion of risk genes which have high prefrontal cortex expression during this prenatal developmental stage tend to have subsequent low prefrontal expression in this region during Late Childhood. In the reciprocal analysis, conditioning on Early Midfetal 1 had little effect on the strong negative correlation between Late Childhood gene expression and schizophrenia association (uncorrected *P*=1.2×10^−6^), implying that there is a greater set of independently associated risk genes with converging expression during this stage.

For both schizophrenia and bipolar disorder, conditioning on Late Childhood did not ablate the relationship between gene expression during Early Infancy and genetic risk (schizophrenia uncorrected *P*=0.0024; bipolar disorder uncorrected *P*=5.9×10^−4^), suggesting that during Early Infancy an independent set of risk genes are differentially expressed.

## DISCUSSION

In this study we have examined the expression of risk genes for schizophrenia and bipolar disorder across development using post mortem gene expression data from the BrainSpan database^39,40^. Our results show a significant bias of schizophrenia risk gene expression in prefrontal cortex to the prenatal period, with a subsequent dynamic change in the collective expression of these risk genes across infancy, childhood and adolescence. In contrast, in bipolar disorder we found no evidence of increased expression of bipolar risk genes in the prenatal period, although we observed a similar pattern of change in bipolar risk gene expression in prefrontal cortex across infancy and childhood to that seen in schizophrenia.

Our finding of increased expression of schizophrenia risk genes in the prenatal period is consistent with previous reports^5,36–38^. In the present study we have examined this enrichment across the full polygenic signal of schizophrenia risk using gene property analyses in MAGMA, suggesting that this bias towards prenatal expression exists across a wide number of schizophrenia associated alleles. Our results show an excess of schizophrenia risk gene expression in the prefrontal cortex during early midfetal development, consistent with findings from key epidemiological studies implicating this period in risk for schizophrenia^3,4,7,8^. This period of development is one of profound neurogenesis and corticogenesis^6^. Subtle alterations in gene expression modulating these processes could represent one form of the early developmental cortical “lesion” proposed by the neurodevelopmental hypothesis of schizophrenia^1,5^. Notably a similar bias of expression of risk associated genes to the prenatal period has also previously been described for autism and intellectual disability^42^, further supporting the view that these disorders can collectively be considered to be neurodevelopmental in origin. However, for bipolar disorder risk genes the elevation in foetal expression was less pronounced and we failed to observe any significant evidence of a prenatal bias in gene expression, consistent with the view that these conditions can be collectively viewed as lying on a spectrum of neurodevelopmental risk, with bipolar disorder characterised by a lower prenatal burden of risk^4,34,35^. These results are also consistent with a recent study showing that polygenic risk for schizophrenia, but not polygenic risk for bipolar disorder, is associated with impairments in early infant neuromotor development^43^.

The transition from late childhood into adolescence also represents a profound period of frontal cortical maturation. In particular, this period is characterised by the elimination (“pruning”) of excitatory synapses in the frontal cortex resulting in changes in excitatory-inhibitory balance and dopaminergic regulation and the emergence of adult patterns of executive functioning^6,24^. It has for some time been hypothesised that risk for schizophrenia and related disorders may be in part mediated through impacts on these late maturational changes in the frontal cortex, which immediately precede the period of peak onset of psychotic disorders^17^. In the present study we find evidence that in both schizophrenia and bipolar disorder there is a profound change in risk gene expression in frontal cortex across the transition from late childhood into adolescence. The pattern in both disorders is of low expression in late childhood which then normalises at the transition into adolescence. Whilst we do not see an overall differential expression of risk genes in adolescence, these data do suggest that there is a dynamic shift in risk gene expression for both schizophrenia and bipolar across this developmental period. It is also notable that a similar pattern of gene expression change is seen in both schizophrenia and bipolar disorder. This finding is consistent with the strong genetic overlap of these conditions and suggests that this overlap derives in part from genes which show dynamic expression changes in the frontal cortex across later childhood and adolescence.

We also found evidence of an increase in risk gene expression for both bipolar disorder and schizophrenia in frontal cortex during early infancy. Early infancy is a critical period of human brain development, but has been less discussed in terms of later risk for psychiatric disorders such as bipolar disorder and schizophrenia than prenatal and adolescent development. Human infancy represents a period of rapid brain maturation including synaptogenesis, myelination and the establishment of patterns of network activation which include frontal regions^44^. These developmental changes sub-serve the emergence of cognitive functions including early language development, attention, working memory and self-regulation^45^, all of which are strongly implicated in neuropsychiatric disorders. Interestingly our conditional analyses show that the genes driving this increase in expression during infancy are not the same as those showing decreased expression in later childhood, implicating differing sets of genes in conferring risk across development. This contrasts with our analysis investigating potential overlaps between genes expressed in prenatal development in schizophrenia and those showing changes across late childhood-adolescence where we find evidence of a substantial overlap in implicated genes through conditional analyses.

Our study has a number of limitations which should be borne in mind. Firstly, the developmental expression data come from a relatively limited group of post-mortem brains, although this dataset is large compared to many comparator datasets and benefits from high quality standards and careful curation^39^. Nevertheless, replication of the current findings in additional samples will be important. In addition, it will be desirable to identify which specific cell types the enrichment signals derive from, given increasing evidence of the involvement of specific key cell types in schizophrenia and related disorders^46,47^. Our study only looks at overall bias of expression of schizophrenia and bipolar associated genes across development and does not implicate specific genes or variants. Future work will be required to substantiate these findings for specific genes and pathways, and it is likely that not all risk genes will conform to the predominant expression patterns identified here^48^. Furthermore, the use of post-mortem gene expression data, whilst powerful, does not necessarily capture all activity-dependent changes in gene expression which may be particularly important in the context of adult plasticity and learning. Finally, it was not possible to generate separate genetic risk profiles for bipolar patients with and without psychotic symptoms, meaning we are not able to say whether genes associated with psychotic bipolar disorder are more strongly expressed in early neurodevelopment, as may be expected from the symptom overlap with schizophrenia^49^.

Overall this work highlights the dynamic changes in expression of risk-associated genes for schizophrenia and bipolar disorder across development. Our results are consistent with the view that there is a significant early neurodevelopmental component to risk for schizophrenia, deriving from processes operating in prenatal development, which is not extensively shared with bipolar disorder. We also provide evidence of a common pattern of changes in risk gene expression for both schizophrenia and bipolar disorder across infancy, childhood and adolescence. Our results show a particularly strong association between risk gene expression in both conditions and prefrontal cortical development, supporting the view that the dynamic and extended development of the human prefrontal cortex is an important substrate for risk for these disorders. Overall our data are consistent with the view that schizophrenia and bipolar disorder lie on a continuum of neurodevelopmental risk^34,35^, with evidence of substantial similarities in risk gene expression across development but a greater burden of risk gene expression in prenatal development in schizophrenia.

## ACKNOWLEDGEMENTS

This work was supported by Medical Research Council (MRC) grant MR/L010305/1, a Wellcome Trust Strategic Award (100202/Z/12/Z) and The Waterloo Foundation ‘Changing Minds’ programme. We thank the Bipolar Disorder Working Group of the Psychiatric Genomics Consortium for providing the bipolar disorder summary statistics used in this study.

## SUPPLEMENTARY INFORMATION

Supplementary Table 1

